# Calculating Biological Module Enrichment or Depletion and Visualizing Data on Large-scale Molecular Maps with ACSNMineR and RNaviCell R packages

**DOI:** 10.1101/064469

**Authors:** Paul Deveau, Emmanuel Barillot, Valentina Boeva, Andrei Zinovyev, Eric Bonnet

## Abstract

Biological pathways or modules represent sets of interactions or functional relationships occurring at the molecular level in living cells. A large body of knowledge on pathways is organized in public databases such as the KEGG, Reactome, or in more specialized repositories, such as the Atlas of Cancer Signaling Network (ACSN). All these open biological databases facilitate analyses, improving our understanding of cellular systems. We hereby describe the R package `ACSNMineR` for calculation of enrichment or depletion of lists of genes of interest in biological pathways. ACSNMineR integrates ACSN molecular pathways, but can use any molecular pathway encoded as a GMT file, for instance sets of genes available in the Molecular Signatures Database (MSigDB). We also present the R package `RNaviCell`, that can be used in conjunction with `ACSNMineR` to visualize different data types on web-based, interactive ACSN maps. We illustrate the functionalities of the two packages with biological data taken from large-scale cancer datasets.

## 1 Introduction

Biological pathways and networks comprise sets of interactions or functional relationships, occurring at the molecular level in living cells [1, 2]. A large body of knowledge on cellular biochemistry is organized in publicly available repositories such as the KEGG database [3], Reactome [4] and MINT [5]. All these biological databases facilitate a large spectrum of analyses, improving our understanding of cellular systems. For instance, it is a very common practice to cross the output of high-throughput experiments, such as mRNA or protein expression levels, with curated biological pathways in order to visualize the changes, analyze their impact on a network and formulate new hypotheses about biological processes. Many biologists and computational biologists establish list of genes of interest (e.g. a list of genes that are differentially expressed between two conditions, such as normal vs disease) and then evaluate if known biological pathways have significant overlap with this list of genes.

We have recently released the Atlas of Cancer Signaling Network (ACSN), a web-based database which describes signaling and regulatory molecular processes that occur in a healthy mammalian cell but that are frequently deregulated during cancerogenesis [6]. The ACSN atlas aims to be a comprehensive description of cancer-related mechanisms retrieved from the most recent literature. The web interface for ACSN is using the NaviCell technology, a software framework dedicated to web-based visualization and navigation for biological pathway maps [7]. This environment is providing an easy navigation of maps through the use of the Google Maps JavaScript library, a community interface with a web blog system, and a comprehensive module for visualization and analysis of high-throughput data [8].

In this article, we describe two R packages related to ACSN analysis and data visualization. The package `ACSNMineR` is designed for the calculation of gene enrichment and depletion in ACSN maps (or any user-defined gene set via the import function), while `RNaviCell` is dedicated to data visualization on ACSN maps. Both packages are available on the Comprehensive R Archive Network (https://cran.r-project.org/web/packages/ACSNMineR/ and https://cran.r-project.org/web/packages/RNaviCell/), and on the GitHub repository (https://github.com/sysbio-curie/ACSNMineR and https://gitlmb.com/sysbio-curie/RNaviCell). For the remainder of this article, we describe the organization of each package and illustrate their capacities with several concrete examples demonstrating their capabilities.

## 2 Packages organization

### 2.1 ACSNMineR

Currently, ACSN maps cover signaling pathways involved in DNA repair, cell cycle, cell survival, cell death, epithelial-to-mesenchymal transition (EMT) and cell motility. Each of these large-scale molecular maps is decomposed in a number of functional modules. The maps themselves are merged into a global ACSN map. Thus the information included in ACSN is organized in three hierarchical levels: a global map, five individual maps, and several functional modules. Each ACSN map covers hundreds of molecular players, biochemical reactions and causal relationships between the molecular players and cellular phenotypes. ACSN represents a large-scale biochemical reaction network of 4,826 reactions involving 2,371 proteins (as of today), and is continuously updated and expanded. We have included the three hierarchical levels in the `ACSNMineR` package, in order to be able to calculate enrichments at all three levels. The calculations are made by counting the number of occurences of gene symbols (HUGO gene names) from a given list of genes of interest in all ACSN maps and modules. Table 1 is detaining the number of gene symbols contained in all the ACSN maps.

**Table 1:** ACSN maps included in the ACSNMineR package. Map: map name, Total: total number of gene symbols (HUGO) used to construct the map, Nb mod.: number of modules, Min: mimimum number of gene symbols in the modules, Max: maximum number of gene symbols in the modules, Mean: average number of gene sybols per module. N.B.: one gene symbol may be present in several modules of the map.

| Map | Total | Nb mod. | Min | Max | Mean |
| --- | --- | --- | --- | --- | --- |
| ACSN global | 2243 | 63 | 2 | 629 | 83 |
| Survival | 1053 | 5 | 208 | 431 | 328 |
| Apoptosis | 667 | 7 | 19 | 382 | 136 |
| EMT & Cell motility | 634 | 9 | 18 | 629 | 136 |
| DNA repair | 345 | 21 | 3 | 171 | 45 |
| Cell cycle | 250 | 25 | 2 | 130 | 19 |

The statistical significance of the counts in the modules is assessed by using either the Fisher exact test [9, 10] or the hypergeometric test, which are equivalent for this purpose [11].

The current ACSN maps are included in the `ACSNMineR` package, as a list of character matrices.

~~~
> length(ACSN_maps)
[1] 6
> names(ACSN_maps)
[1] “Apoptosis” “CellCycle” “DNA_repair” “EMT_motility” “Master”
[6] “Survival
~~~

For each matrix, rows represent a module, with the name of the module in the first column, followed by a description of the module (optional), and then followed by all the gene symbols of the module. The maps will be updated according to every ACSN major release.

The main function of the `ACSNMineR` package is the `enrichment` function, which is calculating over-representation or depletion of genes in the ACSN maps and modules. We have included a small list of 12 Cell Cycle related genes in the package, named genes_test that can be used to test the main enrichment function and to get familiar with its different options.

~~~
> genes_test
[1] “ATM” “ATR” “CHEK2” “CREBBP” “TFDP1” “E2F1” “EP300”
[8] “HDAC1” “KAT2B” “GTF2H1” “GTF2H2” “GTF2H2B”
~~~

The example shown below is the simplest command that can be done to test a gene list for over-representation on the six included ACSN maps. With the list of 12 genes mentionned above and a default p-value cutoff of 0.05, we have a set of 36 maps or modules that are significantly enriched. The results are structured as a data frame with nine columns displaying the module name, the module size, the number of genes from the list in the module, the names of the genes that are present in the module, the size of the reference universe, the number of genes from the list that are present in the universe, the raw p-value, the p-value corrected for multiple testing and the type of test performed. The module field in the results data frame indicate the map name and the module name separated by a column character. If a complete map is significantly enriched or depleted, then only the map name is shown, without any module or column character. For instance, the third line of the results object below concern the E2F1 module of the CellCycle map.

~~~
> library(ACSNMineR)
> results <- enrichment(genes_test)
> dim(results)
[1] 8 9
> results[3,]
                 module module_size nb_genes_in_module
V161 CellCycle:E2F1             19                 12
                                                   genes_in_module
V161 ATM ATR CHEK2 CREBBP TFDP1 E2F1 EP300 HDAC1 KAT2B GTF2H1 GTF2H2 GTF2H2B
     universe_size nb_genes_in_universe      p.value p.value.corrected    test
V161          2237                12 3.735018e-21          2.353061e-19 greater
~~~

The `enrichment` function can take up to eight arguments: the gene list (as a character vector), the list of maps that will be used to calculate enrichment or depletion, the type of statistical test (either the Fisher exact test or the hypergeometric test), the module minimal size for which the calculations will be done, the universe, the p-value threshold and the alternative (“greater” for calculating over-representation, “less” for depletion and “both” for both tests).

Only the gene list is mandatory to call the `enrichment` function, all the other arguments have default values. The maps argument can either be a dataframe imported from a gmt file with the `format_from_gmt` function or a list of dataframes generated by the same procedure. By default, the function uses the ACSN maps previously described. The correction for multiple testing is set by default to use the method of Benjamini & Hochberg, but can be changed to any of the usual correction methods (Bonferroni, Holm, Hochberg, Holm, or Benjamini & Yekutieli [12]), or even disabled. The minimal module size represents the smallest size value of a module that will be used to compute enrichment or depletion. This is meant to remove results of low significance for module of small size. The universe in which the computation is made by default is defined by all the gene symbols contained in the maps. All the genes that were given as input and that are not present on the maps will be discarded. To keep all genes, the user can change the universe to `HUGO`, and in that case, the complete list of HUGO gene symbols will be used as the reference (> 39,000 genes). The threshold corresponds to the maximal value of the corrected p-value (unless the user chose not to correct for multiple testing) that will be displayed in the result table.

It may be of interest to compare enrichment of pathways in different cohorts or experiments. For example, enrichment of highly expressed pathways can reveal differences between two cancer types or two cell lines. To facilitate such comparisons, `ACSNMineR` provides a `multisample_enrichmerLt` function. It relies on the enrichment function but takes a list of character vector genes. The name of each element of the list will be assumed to be the name of the sample for further analysis. Most of the arguments given to `multisample_enrichment` are the same as the ones passed to `enrichment`. However, the `cohort_threshold` is designed to filter out modules which would not pass the significance threshold in all samples.

Finally, to facilitate visualization of results, `ACSNMineR` integrates a representation function based on `ggplot2` syntax [13]. It allows representation of results from `enrichment` or `multisample_enrichment` with a limited number of parameters. Two types of display are available: heat-map tiles or bars. For multiple samples using a barplot representation, the number of rows used can be provided, otherwise all plots will be on the same row. For the heatmap, the color of the non-significant modules, and boundaries of the gradient for significant values can also be tuned.

We previously computed the p-value of the `genes_test` list with default parameters. The number of modules which have a p-value below 0.05 was 36, that can be compared to the 49 obtained without correction with the simple command shown below (some of the results are displayed in table 2).

~~~
.enrichment(genes_test,correction_multitest = FALSE)
~~~

**Table 2:**
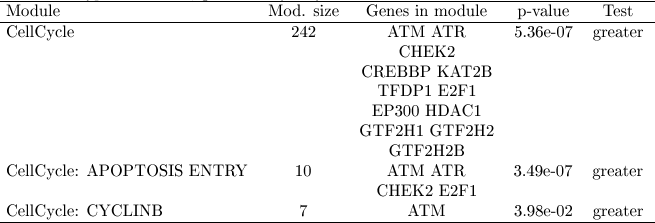
Overview of the results from enrichment analysis without correction. Module: name of the module. Mod. size: size of the module. Genes in module: genes from input which are found in the module. p-value: uncorrected p-value. Test: null hypothesis used, greater is synonym of enrichment.

We can now plot the first six rows of the results obtained for corrected and uncorrected fisher test with heatmap format (Figure 1) or barplot (Figure 2) with the following commands:

~~~
# heatmap

represent_enrichment(enrichment = list(Corrected = results[1:6,],
Uncorrected = results_uncorrected[1:6,]),
                               plot = “heatmap”, scale = “reverselog”,
                               low = “steelblue”, high =“white”, na.value = “grey”)

# barplot

represent_enrichment(enrichment = list(Corrected = results[1:6,],
                Uncorrected = results_uncorrected[1:6,]),
                 plot = “heatmap”, scale = “reverselog”,
                 nrow = 1)
~~~

**Figure 1:**
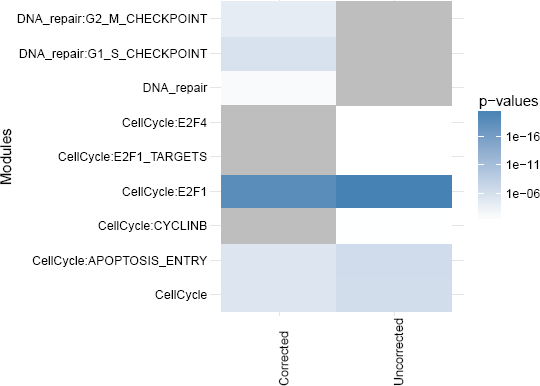
Representation of the enriched modules (first six rows), with either Bonferroni correction or no correction. Grey tiles means that the data is not available for this module in this sample. P-values of low significance are in white, whereas p-values of high significance are represented in blue.

**Figure 2:**
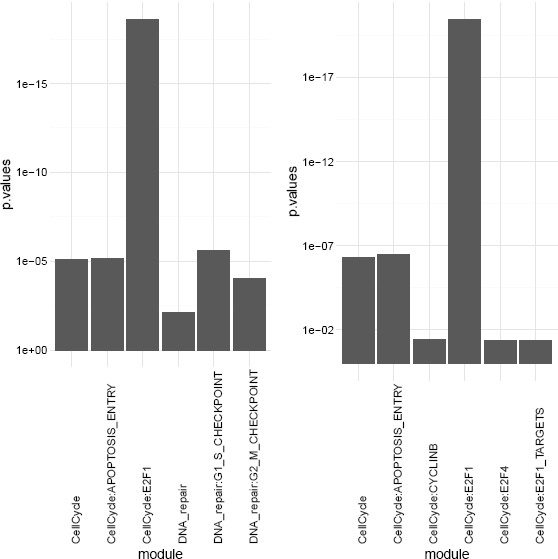
Representation of the enriched modules (first six rows), with either Bonferroni correction (left) or no correction (right). The modules are on the X axis and the p-values are on the Y axis.

### 2.2 RNaviCell

The NaviCell Web Service provides a server mode, which allows automating visualization tasks and retrieving data from molecular maps via RESTful (standard http/https) calls. Bindings to different programming languages are provided in order to facilitate the development of data visualization workflows and third-party applications [8]. RNaviCell is the R binding to the NaviCell Web Service. It is implemented as a standard R package, using the R object-oriented framework known as Reference Classes [14]. Most of the work done by the user using graphical point-and-click operations on the NaviCell web interface are encoded as functions in the library encapsulating http calls to the server with appropriate parameters and data. Calls to the NaviCell server are performed using the library RCurl [15], while data encoding/decoding in JSON format is performed with the RJSONIO library [16].

Once the RNaviCell library is installed and loaded, the first step is to create a NaviCell object and launch the browser session. This will automatically create a unique session ID with the NaviCell server. Once the session is established, various functions can be called to send data to the web session, set graphical options, visualize data on a map or get data from the map. There are 125 functions available in the current version of RNaviCell. All of them are described with their different options in the RNaviCell documentation, and we provide a tutorial on the GitHub repository wiki (https://github.com/sysbio-curie/RNaviCell/wiki/Tutorial).

In the simple example detailed below, we create a NaviCell session, then load an expression data set from a local (tab-delimited) file. The data represent gene expression measured in a prostate cancer cell line resistant to hormonal treatment (agressive), and is taken from the Cell Line Encyclopedia project [17]. We visualize the data values on the Cell Cycle map (the default map), using heat maps. With this visualization mode, gene expression values are represented as a color gradient (green to red) in squares positioned next to the entities where the gene has been mapped (Figure 3).

~~~
# a short RNaviCell script example

# load RNaviCell library

library(RNaviCell)

# create a NaviCell object and launch a server session
# this will automatically open a browser on the client

navicell <- NaviCell()
navicell$launchBrowser()

# import a gene expression matrix and
# send the data to the NaviCell server
# NB: the data_matrix object is a regular R matrix

data_matrix <- navicell$readDatatable(‘DU145_data.txt’)
navicell$importDatatable(“mRNA expression data”, “DU145”, data_matrix)

# set data set and sample for heat map representation

navicell$heatmapEditorSelectSample(‘0’,‘data’)
navicell$heatmapEditorSelectDatatable(‘0’,‘DU145’)
navicell$heatmapEditorApply()
~~~

**Figure 3:**
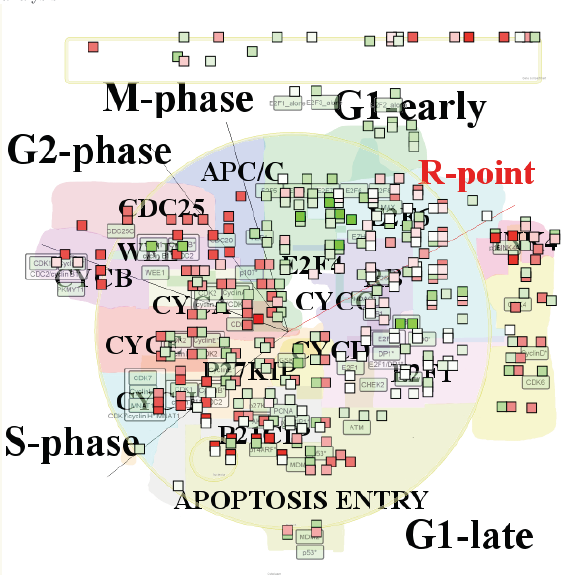
Gene expression values from a prostate cancer cell line visualized on the cell cycle map as heat map plots. The figure is a screenshot of the NaviCell map browser, with the map set at the top (the less detailed) zoom level. The essential phases of the cell cycle are indicated on the map (G1/S/G2/M). Note that on the web browser the map is interactive and the user can zoom in and out, change the graphical parameters, import additional data and perform functional analysis.

## 3 Case studies

### 3.1 Analysis of breast cancer expression data

In a study published in 2008, Schmidt and colleagues analyzed gene expression patterns of 200 breast cancer patients not treated by systemic therapy after surgery using discovery approach to reveal additional prognostic motifs [18]. Estrogen receptor (ER) expression and proliferative activity of breast carcinomas are well-known and described prognostic markers. Patients with ER-positive carcinomas have a better prognosis than those with ER-negative carcinomas, and rapidly proliferating carcinomas have an adverse prognosis. Knowledge about the molecular mechanisms involved in the processes of estrogen-dependent tumor growth and proliferative activity has led to the successful development of therapeutic concepts, such as antiendocrine and cytotoxic chemotherapy.

The dataset corresponding to this study is available as a Bioconductor package. The code shown below is creating a list of differentially expressed genes between ER positive and ER negative samples, and calculates the enrichment in ACSN maps from this list of genes. As seen in Table 3, there is one map (Cell Cycle) and six modules (belonging to the Cell Cycle, DNA repair and EMT motility maps) enriched.

~~~
# load all necessary packages
library(breastCancerMAINZ)
library(Biobase)
library(limma)
library(ACSNMineR)
library(hgu133a.db)
library(RNaviCell)

# load data and extract expression and phenotype data
data(mainz)
eset <- exprs(mainz)
pdat <- pData(mainz)

# Create list of genes differentially expressed between ER positive and
# ER negative samples using moderated t-test statistics
design <- model.matrix(~factor(pdat$er == ‘1’))
lmFit(eset, design) -> fit
eBayes(fit) -> ebayes
toptable(ebayes, coef=2,n=25000) -> tt
which(tt$adj < 0.05) -> selection
rownames(tt[selection,]) -> probe_list
mget(probe_list, env=hgu133aSYMBOL) -> symbol_list
symbol_list <- as.character(symbol_list)

# calculate enrichment in ACSN maps

enrichment(symbol_list) -> results

dim(results)
[1] 7 9
~~~

**Table 3:** ACSN maps enrichment for genes differentially expressed between ER positive and ER negative samples in breast cancer. Module: name of the module. Mod. size: size of the module. Nb genes: number of genes from input which are found in the module, pval: raw p-value. Cor. pval: corrected p-value.

| Module | Mod. size | Nb genes | pval | Cor. pval |
| --- | --- | --- | --- | --- |
| CellCycle | 242 | 20 | 1.70e-03 | 0.021 |
| CellCycle:E2F1_TARGETS | 130 | 14 | 6.26e-04 | 0.009 |
| CellCycle:E2F2_TARGETS | 35 | 8 | 4.70e-05 | 0.009 |
| CellCycle:E2F3_TARGETS | 51 | 11 | 3.82e-06 | 0.0002 |
| CellCycle:E2F4_TARGETS | 100 | 15 | 9.99e-06 | 0.0003 |
| DNA_repair:SPINDLE_CHECKPOINT | 28 | 5 | 3.82e-03 | 0.04 |
| EMT_motility:ECM | 147 | 13 | 5.09e-03 | 0.045 |

The Molecular Signatures Database (MSigDB) is one of the most widely used repository of well-annotated gene sets representing the universe of biological processes [19]. We downloaded the canonical pathways set, counting more than 1,300 gene sets representing canonical pathways compiled by domain experts. The dataset is encoded with the ‘gmt’ format, and can be imported within ACSNMineR with the `format_from_gmt` function. We calculate the enrichment for the breast cancer differentially expressed gene list, simply specifying the MSigDB data we just imported as the maps option. Table 4 is displaying the pathways having a corrected p-value < 0.05. The prefix is indicating the database source, so we see that we have pathways from the KEGG, Reactome and PID databases. Consistent with our previous results, most of the enriched pathways are related to the cell cycle regulation.

~~~
# Import MSigDB canonical pathways and calculate enrichment on this database

mtsig <- format_from_gmt(‘c2.cp.v5.0.symbols.gmt’)
enrichment(symbol_list, maps = mtsig)
~~~

**Table 4:**
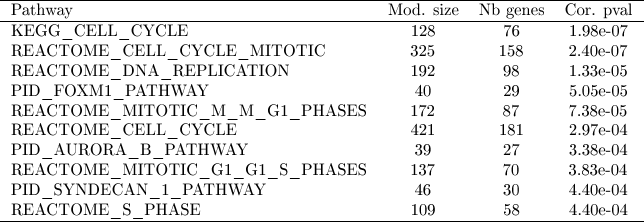
MSigDB canonical pathway database enrichment for genes differentially expressed between ER positive and ER negative samples in breast cancer. Module: name of the module. Mod. size: size of the module. Nb genes: number of genes from input which are found in the module. Cor. pval: corrected p-value.

At last, we visualize the mean expression values for ER negative samples for all genes differentially expressed on the ACSN master (global) map using `ACSNMineR` commands to create heatmaps.

~~~
# Select ER negative samples and calculate mean expression values

apply(eset[probe_list,pdat$er == 0],1,mean) -> er_minus_mean
names(er_minus_mean) <- symbol_list
er_minus_mean <- as.matrix(er_minus_mean)
colnames(er_minus_mean) <- c(‘exp’)

# create a NaviCell session, import the expression matrix on the map and create
# heatmaps to represent the data points.

navicell <- NaviCell()
~~~

~~~
navicell$proxy_url <- “https://acsn.curie.fr/cgi-bin/nv_proxy.php”
navicell$map_url <- “https://acsn.curie.fr/navicell/maps/acsn/master/index.php”

navicell$launchBrowser()
navicell$importDatatable(“mRNA expression data”, “GBM_exp”, er_minus_mean)
navicell$heatmapEditorSelectSample(‘0’,‘exp’)
navicell$heatmapEditorSelectDatatable(‘0’,‘GBM_exp’)
navicell$heatmapEditorApply()
~~~

The Figure 4 is displaying the map for gene that having a corrected p-value < 0.05 and a log fold-change > 1. we can see that the genes are concentrated in the regions of the map corresponding to the cell cycle, cell motility, apoptosis and survival.

### 3.2 Analysis of glioblastoma mutation frequencies

Recent years have witnessed a dramatic increase in new technologies for interrogating the activity levels of various cellular components on a genome-wide scale, including genomic, epigenomic, transcriptomic, and proteomic information [20]. Integrating these heterogeneous datasets provides more biological insights than performing separate analyses. For instance, international consortia such as The Cancer Genome Atlas (TCGA) have launched large-scale initiatives to characterize multiple types of cancer at different levels on hundreds of samples. These integrative studies have already led to the identification of novel cancer genes [21].

Malignant gliomas, the most common subtype of primary brain tumors, are aggressive, highly invasive, and neurologically destructive tumors considered to be among the deadliest of human cancers. In its most aggressive manifestation, glioblastoma (GBM), median survival ranges from 9 to 12 months, despite maximum treatment efforts [22]. In this study we have analyzed whole-genome mutation data generated by the TCGA project on hundreds of patients. More specifically, we parsed the MAF (Mutation Annotation Format) GBM files produced by different sequencing centers to count and calculate gene mutation frequencies. We kept the mutations having a status likely to disturb the target protein’s function (i.e, Frame_Shift_Del, Nonstop_Mutation, In_Frame_Del, In_Frame_Ins, Missense_Mutation, Nonsense_Mutation, Splice_Site, Translation_Start_Site). In total, we collected mutations for more than 13,000 genes in a total of 379 mutated samples. In order to retain the most frequently mutated genes, we calculated frequencies across all mutated samples, and kept genes having a frequency greater than 1% (3,293 genes). We further labelled genes having a frequency greater than 1% and less than 5% as “1” and genes highly mutated (frequency higher than 5%) as “2”.

**Figure 4:**
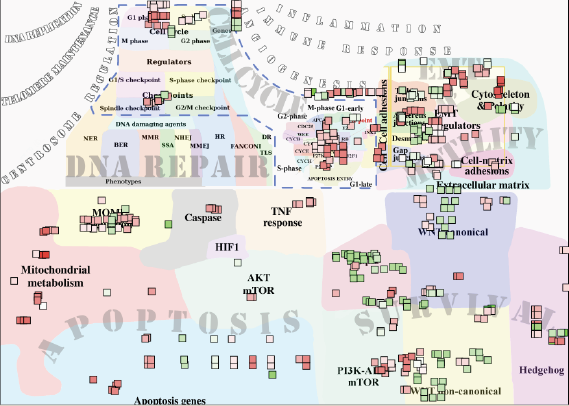
Mean expression values for ER negative differentially expressed genes in breast cancer visualized as heatmaps on the ACSN master map.

We loaded the data as a matrix in R and calculated the enrichment in ACSN maps with the ACSNMineR function `enrichment`. The results are displayed in table 5. There are 6 modules significantly enriched in the DNA repair and EMT motility maps. Cell matrix adhesions and ECM (extra cellular matrix), part of the EMT motility map, are the modules with highest significance. The EMT motility map is significantly enriched at the global map level (second line in the table).

**Table 5:** ACSN maps enrichment for frequently mutated glioblastoma genes. Module: name of the module. Mod. size: size of the module. Nb genes: number of genes from input which are found in the module. Cor. pval: corrected p-value.

| module | Mod. size | Nb genes | Cor. pval |
| --- | --- | --- | --- |
| DNA_repair:S_PHASE_CHECKPOINT | 45 | 19 | 0.008 |
| EMT_motility | 635 | 181 | 0.0002 |
| EMT_motility:CELL_MATRIX_ADHESIONS | 73 | 45 | 3.73e-12 |
| EMT_motility:CYTOSKELETON_POLARITY | 154 | 47 | 0.022 |
| EMT_motility:DESMOSOMES | 29 | 15 | 0.002 |
| EMT_motility:ECM | 147 | 69 | 9.77e-11 |
| EMT_motility:EMT_REGULATORS | 629 | 178 | 0.0002 |

Visualization of the list of glioblastoma mutated genes is shown on figure 5. This figure was generated with the ACSNMineR commands detailed below. Results of the enrichment test correlate well with the visualization on the map, with a high density of low and high frequency mutated genes in the EMT motility and DNA repair regions (maps) of the global ACSN map. Although they are not statistically significant, quite high densities can also be seen in other regions of the map.

~~~
library(RNaviCell)

# Create a NaviCell object, point it to the ACSN master map and launch
# a session.

navicell <- NaviCell()
navicell$proxy_url <- “https://acsn.curie.fr/cgi-bin/nv_proxy.php”
navicell$map_url <- “https://acsn.curie.fr/navicell/maps/acsn/master/index.php”
navicell$launchBrowser()

# Read the GBM data file and import it into the session.

mat <- navicell$readDatatable(‘gbm.txt’)
navicell$importDatatable(“Mutation data”, “GBM”, mat)

# set datatable and sample names for the glyph editor

navicell$drawingConfigSelectGlyph(1, TRUE)
navicell$glyphEditorSelectSample(1, “categ”)
navicell$glyphEditorSelectShapeDatatable(1, “GBM”)
navicell$glyphEditorSelectColorDatatable(1, “GBM”)
navicell$glyphEditorSelectSizeDatatable(1, “GBM”)
navicell$glyphEditorApply(1)

# set color, shape and size parameters for glyphs

navicell$unorderedConfigSetDiscreteShape(“GBM”, “sample”, 0, 1)
navicell$unorderedConfigSetDiscreteShape(“GBM”, “sample”, 1, 5)
navicell$unorderedConfigApply(“GBM”, “shape”)

navicell$unorderedConfigSetDiscreteColor(“GBM”, “sample”, 0, “398BC3”)
navicell$unorderedConfigSetDiscreteColor(“GBM”, “sample”, 1, “CC5746”)
navicell$unorderedConfigApply(“GBM”, “color”)

navicell$unorderedConfigSetDiscreteSize(“GBM”, “sample”, 0, 4)
navicell$unorderedConfigSetDiscreteSize(“GBM”, “sample”, 1, 14)

navicell$unorderedConfigApply(“GBM”, “size”)
~~~

**Figure 5:**
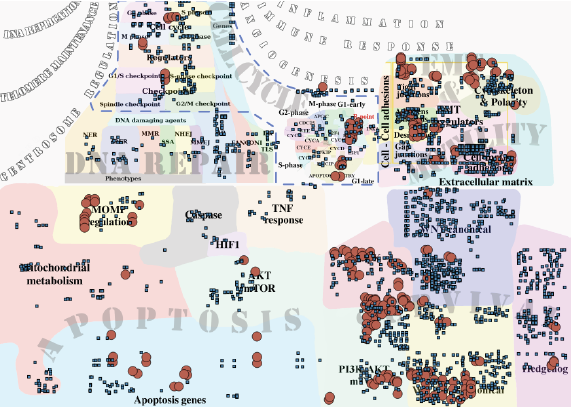
Glioblastoma gene mutation frequency categories represented as glyphs on the ACSN global cancer map. High frequency mutated genes are pictured as large red circles, while low frequency mutated genes are depicted as small blue squares.

## 4 Summary and perspectives

In this work, we presented the `R` package `ACSNMineR`, a novel package for the calculation of p-values for enrichment or depletion of genes in biological pathways. The package includes the six large-scale molecular maps and 55 functional modules of the Atlas of Cancer Signaling Network (ACSN). Enrichment can be calculated for those maps and modules with several options to play with, but can also be calculated for other databases of molecular pathways, that can be imported from GMT formated files.

We also describe in this work the `RNaviCell` package, a `R` package convenient to use with `ACSNMineR`. This package is dedicated to create web-based and interactive data visualization on ACSN maps. Users can use this tools to represent genes of interest that have been shown to be related to the maps by calculating enrichment with the `ACSNMineR`. Creating maps with the graphical user interface of the ACSN website can be a tedious task if the user has multiple samples or gene lists, and wants to compare their representations on ACSN maps. The `RNaviCell` package can be used to automate the process of creating the graphical representations automatically.

We have shown how the packages `ACSNMineR` and `RNaviCell` can be combined to analyze expression data from breast cancer samples, and also to analyze the frequency of mutated genes in glioblastoma cancer samples.

Of course, `ACSNMineR` is not the only `R` package for enrichment calculations. For instance, `GOstats` [23] is probably one of the first packages that was created to calculate enrichment for Gene Ontology categories. `GOstats` can also be used to calculate enrichment for other biological pathways categories, such as KEGG pathways (by using an instance of the class `KEGGHyperGParams`) or PFAM protein families (using `PFAMHyperGParams`). However, its usage might not be as straightforward as `ACSNMineR`, and it does not seem possible to test user-defined biological pathways. Furthermore, other authors have pointed out that the KEGG database used by this package has not been updated since 2012. `clusterProfiler` is a recent R package released for enrichment analysis of Gene Ontology and KEGG with either hypergeometric test or Gene Set Enrichment Analysis (GSEA) [24]. Via other packages, support for analysis of Disease Ontology and Reactome Pathways is possible. Interestingly, this package also offers the possibility to import user-defined gene set, through tab-delimited pairwise definition files.

In order to improve `ACSNMineR`, we may in the near future try to improve the speed of calculation, which might be a problem if a very large number of samples or experiments have to be analyzed rapidly. For instance, we could use the `foreach` and `\%dopar\%`, operator to parallelize the most computationally demanding operations. It could also be useful to implement more sensitive methods of gene set enrichment measures, such as the Gene Set Enrichment Analysis (GSEA) method.

## 5 Acknowledgements

This work was supported by a grant “Projet Incitatif et Collaboratif Computational Systems Biology Approach for Cancer” from Institut Curie. The authors would like to thank Pierre Gestraud for his comments on early versions of the `ACSNMineR` package and Eric Viara for guidance and assistance on the development of the `RNaviCell` package.

